# Spontaneous activation of cortical somatosensory networks depresses their excitability in preterm human neonates

**DOI:** 10.1101/2022.12.08.519675

**Authors:** Kimberley Whitehead, Maria Pureza Laudiano-Dray, Judith Meek, Sofia Olhede, Lorenzo Fabrizi

## Abstract

In the developing cortex of preterm human infants, neuronal activity is discontinuous – characterized by sudden, high-amplitude bursts that interrupt periods of quiet background activity. While the functional significance of these bursts is well established, the underlying cause remains unclear.

We propose that this burst–quiescence pattern arises from a temporary “refractoriness” in cortical networks following spontaneous activation. To investigate this, we examined whether spontaneous activity in sensory networks reduces their excitability by assessing how ongoing brain activity influences responses to external sensory stimuli.

We recorded electroencephalographic (EEG) responses to tactile stimulation of the hands and feet in 35 preterm infants, with a median post-menstrual age of 32 weeks. This stimulation triggered increases in wideband cortical power, showing two distinct peaks: one in the delta range and another in the alpha-beta range. Delta-band activity is widespread across the scalp, while the faster alpha-beta responses were confined to somatotopically specific regions.

Importantly, we found that when the baseline activity shared similar spectral and spatial characteristics with the evoked somatosensory response, the magnitude of the evoked response was reduced. This suggests that spontaneous events transiently engage and saturate both widespread (tangential) and localized (columnar) cortical circuits.

As a result, the same cortical regions become temporarily less responsive—a form of refractoriness—preventing immediate reactivation. This mechanism may explain the cyclical pattern of bursting and quiescence observed in the preterm brain.

**Significance Statement:** It is well known that the preterm human brain exhibits a characteristic alternation between high-amplitude activity and quiescence, yet the underlying mechanism remains unclear. Drawing inspiration from developmental neuroscience in animal models, the present study provides the first potential neurobiological explanation for this rhythmic pattern. Using EEG recordings and a somatosensory stimulation paradigm, it demonstrates that preterm cortical bursts induce a refractory period during which external stimuli fail to elicit a response - revealing an intrinsic, activity-dependent depression mechanism. By presenting fundamental novel insights into the neurobiology of preterm cortical activity, this work has broad implications, offering both basic neuroscientists and clinical specialists new understanding of the developmental origins of sensory processing and neonatal EEG patterns linked to later neurobehavioural outcomes.

## Introduction

Organised spontaneous activity across the central nervous system is a characterising feature of developing vertebrate neuronal circuits. This activity is necessary for initial development of neuronal ensembles and, in the cortex, engages columnar and extended networks, establishing the template of the mature functional architecture before the arrival of sensory signals (Molnár et al., 2020; Martini et al., 2021). This spontaneous activity is characteristically discontinuous where sudden high amplitude bursts of synchronous neuronal activity interrupt periods of quiescent background (Khazipov and Luhmann, 2006). This cyclical pattern can be explained by an excitatory recurrent network that encounters activity-dependent depression (Tabak et al., 2000; Kirmse and Zhang, 2022; Dutta et al., 2023). Indeed, spontaneous activity transiently depresses synaptic responsiveness to incoming stimuli in embryonic spinal cord preparations (“refractoriness”), possibly because of transmitter depletion and ligand-induced changes in postsynaptic currents, which slowly recovers in the inter-burst interval (Fedirchuk et al., 1999). This refractory period plays a key role in defining the propagation domains of spontaneous retinal waves (Feller et al., 1996; Godfrey and Swindale, 2007) and is related to after-hyperpolarizations of amacrine cells (Zheng et al., 2006) which also occur in spinal neurons (O’Donovan, 1999). In regards to its function, the episodic nature of activity bouts is thought to provide a high signal-to-noise ratio to the circuit, which could enhance synaptic efficacy in developing networks (O’Donovan, 1999; Tiriac et al., 2014).

Early cortical activity in humans is also highly discontinuous. The preterm electroencephalograph (EEG) is characterised by high voltage transient events (delta brushes and temporal theta) separated by low amplitude inter-burst intervals until about 35 weeks post-menstrual age (Bourel-Ponchel et al., 2021; Wallois et al., 2021). These spontaneous activity bursts occur with varying topographical configurations (Arichi et al., 2017) which sometimes resemble those evoked by external stimulation (e.g. for tactile stimulation: Fig. 3 in (Whitehead et al., 2016)). The diagnostic and prognostic importance of this characteristic pattern of activity is well established (Watanabe et al., 1999; Guérit et al., 2009; Whitehead et al., 2016), however, its aetiology, i.e. why the preterm EEG is discontinuous, is poorly understood. Here, we hypothesise that, as in other species, the preterm EEG is highly discontinuous because of “refractoriness” of cortical networks following spontaneous activation. To test this hypothesis, we assessed whether spontaneous activity in sensory networks depressed their excitability, by measuring the impact of ongoing activity on the response to an external sensory stimulus.

## Materials and Methods

### Subjects

Thirty-five preterm infants were recruited for this study from the neonatal wards at the Elizabeth Garrett Anderson wing of University College London Hospitals (Table 1). Ethical approval was obtained from the NHS Research Ethics Committee, and informed written parental consent was obtained prior to each study. No neonates were acutely unwell at the time of study, required mechanical ventilation, or were receiving analgesics or anti-seizure medication. Exclusion criteria included a diagnosed chromosomal abnormality, high-grade germinal matrix-intraventricular haemorrhage (grade III or above), or birth weight below the 2^nd^ centile. All EEGs were assessed as normal for postmenstrual age by a clinical neurophysiologist (KW) (Tsuchida et al., 2013).

**Table 1.** Demographics of the sample.

|  | Total |
| --- | --- |
| No. of neonates | 35 |
| No. of stimulation trains | 101 (48 hands, 53 feet) |
| Median (range) PMA at time of study (weeks+days) | 32+6 (28+2-35+5) |
| Median (range) PNA at study (days) | 7 (2-33) |
| Sex (% female) | 40% |
| Multiple gestation neonates | 31% |
| Median (range) birth weight (g) | 1575 (785-2440) |

PNA indicates postnatal age; PMA indicates postmenstrual age: gestational age at birth plus PNA; Preterm = <37 weeks gestational age at birth.

### EEG recording set-up

Up to nineteen recording electrodes (disposable Ag/AgCl cup electrodes) were positioned according to the modified international 10/10 electrode placement system at F3, F4, FCz, Cz, CPz, C3, C4, CP3, CP4, F7, F8, T7, T8, P7, P8, TP9, TP10, O1 and O2. A reduced number of electrodes were applied if the infant became unsettled during set-up (median 18; range 11-19). The reference electrode was placed at Fz and the ground electrode was placed at FC5/6. Target impedance of electrodes was <10 kΩ (André et al., 2010). EEG was recorded with a direct current (DC)-coupled amplifier from DC-800Hz using a Neuroscan (Scan 4.3) SynAmps2 EEG/EP recording system. Signals were digitized with a sampling rate of 2 kHz and a resolution of 24 bit.

### Sensory stimulation

Tactile stimuli were delivered by tapping the infants’ hands and feet while naturally asleep in the cot or held by their parents, with a hand-held tendon hammer (Supplementary Video 1) (Worley et al., 2012). A train of somatosensory stimuli was delivered to each limb. The interstimulus interval was large, variable, and self-paced by the experimenter (usually 8–20 s). In case the infant moved, the tap was delayed by several seconds to avoid potential modulation of the somatosensory response by the movement (Saby et al., 2016) and allow movement artefacts to resolve. The sequence in which the limbs were stimulated varied across subjects. We attempted to stimulate every limb at every test occasion, but this was not always possible, for example when a cannula was present, or if the infant became unsettled. This resulted in a total of 101 stimulation trains (i.e. stimulated limbs) of 2-48 taps (mean 15).

### EEG pre-processing

Data pre-processing was carried out using EEGLAB v.13 (Swartz Center for Computational Neuroscience) and custom written MATLAB functions. Continuous EEG data were filtered with a comb filter consisting of notch filters at 50 Hz and its harmonics (100 and 150 Hz) to remove mains noise. Each notch filter was a band-stop 4th order zero-phase shift Butterworth filter with a bandwidth of 4 Hz. Continuous EEG data were then epoched from −3 to +3 seconds around the stimulus. This led to 804 foot-stimulus epochs and 700 hand-stimulus epochs. Bad channels (poor contact with the scalp) were removed from 13 datasets (median: 1 channel, range: 1-3 channels) and then estimated with spherical interpolation as implemented in EEGLAB. EEG recordings following stimulation of the left hemi-body were ‘side-swapped’ so that recordings contralateral to the stimulation site appear on the ‘same side’ of the scalp. There are no differences between somatosensory responses to right vs. left body stimulation in preterm infants (Leikos et al., 2020).

### Analysis of stimulus-evoked EEG power changes

We first defined the functional engagement of somatosensory cortical networks as the spectro-spatial characteristics of the EEG power changes elicited by tactile stimulation of hands and feet across the whole scalp. To do that, we compared the median power periodogram (0.2-40 Hz) of the 3-seconds post-stimulus segment to that of the 3-seconds baseline at every channel. Each segment was tapered with the first discrete prolate spheroidal sequence before calculating the individual power periodogram. We then took the median power periodogram across all trials for the baseline 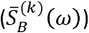 and post-stimulus 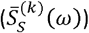 segments at each channel *k* (where *ω*denotes frequency). To assess whether there was a significant difference between these quantities (i.e. whether there was a significant change in the median power periodogram following stimulation) we computed the ratio 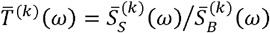 and assessed its significance as previously described (Fabrizi et al., 2016). Significance threshold was set at *α* = 0.05, however, because this test was conducted at any frequency *ω* of the frequency grid, we used false discovery rates to correct for multiple comparisons. Moreover, we also used Bonferroni correction to account for testing at multiple channels (n = 19). The p-values in Fig. 1e and 2e incorporate these adjustments. We then plotted the scalp distributions of these changes in the slow-delta (0.2-2 Hz), fast-delta (2-4 Hz), theta (4-8 Hz), alpha-low beta (8-20 Hz) and high beta-gamma (20-40 Hz) bands on a 3D head mesh for illustration purposes.

**Figure 1.**
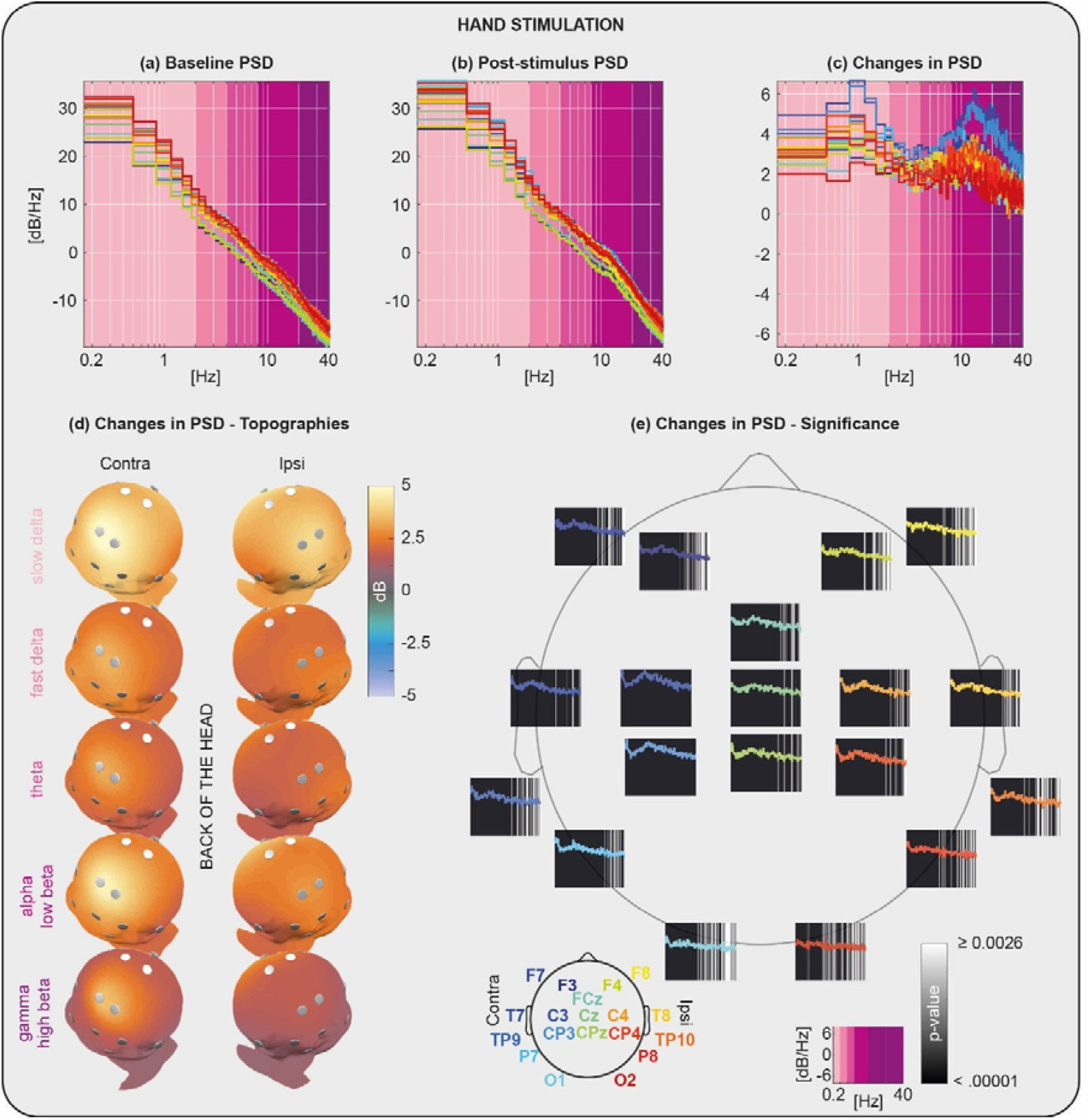
Spectro-spatial EEG power changes associated with tactile stimulation of the hands. Power spectral density (PSD) at baseline **(a)** and following stimulation **(b)** at each channel. PSD of changes elicited by the stimulation **(c)** and 3D maps of their scalp distribution in the slow-delta (0.2-2 Hz), fast-delta (2-4 Hz), theta (4-8 Hz), alpha-low beta (8-20 Hz) and high beta-gamma (20-40 Hz) band **(d)**. Statistical significance of the power changes at each channel **(e)**, note that the changes are the same as those in panel (c) but on a linear frequency axis to allow the display of higher frequencies. P-value limits relate to adjustment for multiple comparisons.

### Analysis of spontaneous EEG spectro-spatial profile effect on stimulus-evoked changes

We then wanted to test whether the spontaneous activation of somatosensory cortical networks influenced their stimulus-driven recruitment. As we had no control over the ongoing activity at the time of stimulation, we had to assess whether the somatosensory networks happened to be active before each of our stimuli. To do that we compared the spectro-spatial profile of the EEG power preceding each stimulus with that of the median EEG power changes elicited by tactile stimulation (somatosensory-evoked template). A close match between them would imply that the somatosensory cortical networks were spontaneously active before the stimulus. To assess this, we used a linear regression model of the form:

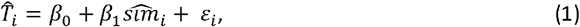

where 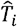 represents a summary measure of observed change in EEG power elicited by stimulus *i* (stimulus response) and 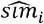 the similarity between the EEG power preceding the stimulus and that of the average median EEG power changes elicited by tactile stimulation. We now define these quantities.

We first estimated the overall power (energy) content 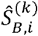 and 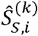 preceding and following each stimulus *i* and the stimulus-related change 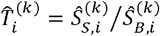 at every electrode *k*, where:

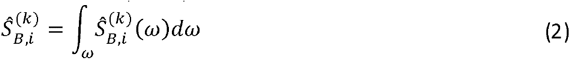

And

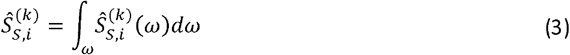

where 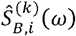 and 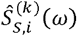 are the EEG spectral estimates preceding and following each stimulus *i*. 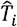 and 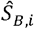 were then calculate as the Euclidean norm of these power vectors in channel space and converted to decibels as:

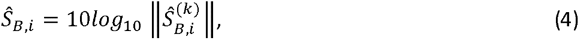

And

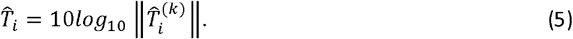

To calculate 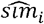, we compared the topographical distribution of the EEG power preceding each stimulus with that of the average median EEG power change elicited by tactile stimulation at five EEG bands (slow-delta, 0.2-2 Hz, Ω_1_; fast-delta, 2-4 Hz, Ω_2_; theta, 4-8 Hz, Ω_3_; alpha-low beta, 8-20 Hz, Ω_4_ and high beta-gamma, 20-40 Hz, Ω_5_). For each band, we defined the average median EEG power change elicited by tactile stimulation (this can be interpreted as the somatosensory cortical networks activation) as:

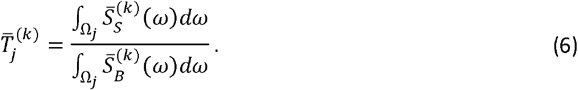

Where we defined:

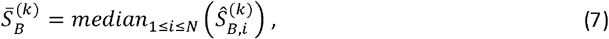

and

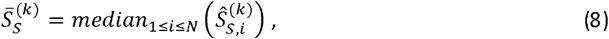

With *N* being the total number of trials.

We also defined the EEG power at each frequency band *j* preceding each stimulus *i* at electrode k as:

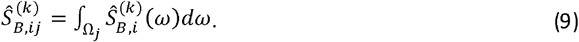

We then calculated the cosine similarity between the topographies of (6) and (9) for each band *j*:

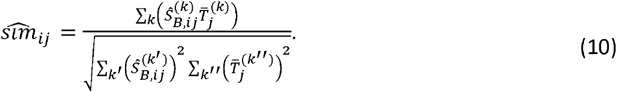

and sum (10) across all bands Ω_J_ to obtain a 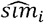 which ranges between 0-5, where 0 is no similarity between topographies and 5 is an exact match:

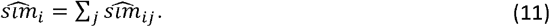

Post-menstrual age (*PMA*) is the primary source of inter-individual variance in the neonatal EEG (Whitehead et al., 2018a). Since in this study we were not interested in the effect of *PMA*, before fitting model (1) we first regressed, 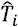 and 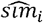 on *PMA*, essentially removing *PMA* effect from all the variables.

In eqn (1) we perform linear regression between 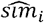 and 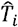 From Fig. 3 we can deduce that the variance of 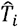 is approximately constant over *i*. It is still not quite in the usual regression framework as 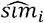 is a random variable. However as it has been averaged over frequency bands Ω_j_ (this only done for 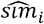 and not for 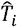), frequencies and channels, we assume that the random variation in 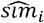 is negligible relative to the random variation in ε _*i*_For this reason, we can perform hypothesis testing for the model in eqn (1), as in the usual linear regression setting with normal theory assumption (NTA, as the multiple summations will make 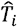 near Gaussian) and second order assumption (SOA).

## Data Accessibility

The data described will be made accessible on Open Science Framework at publication, alongside our existing (separate) open-access neonatal EEG dataset on that portal (Whitehead and Fabrizi, 2024).

## Code Accessibility

Analysis software will either be posted alongside the data (see above) or accessible upon request.

## Results

### Somatosensory-evoked slow neuronal activity is widespread across the scalp, while fast activity engages local somatotopically specific areas

We first defined the spectro-spatial characteristics of somatosensory-evoked activity in 35 preterm infants of median 32 weeks post-menstrual age (Table 1). Tactile stimulation of hands and feet evoked wideband power increases across all scalp EEG channels (Fig. 1 and 2; channel locations are plotted in Fig. 1e and 2e). There were two distinct peaks within the slow-delta band (6.7 and 3.7 dB at 0.9 Hz for hands and feet) and alpha-beta band (6.3 and 4.6 dB at 13 Hz for hands and feet, Fig. 1c and 2c). The topographic distribution of these power changes was frequency dependent. Significant power changes (compared to baseline) at slower frequencies (< 20 Hz) were widespread across the scalp, while changes at faster frequencies were focussed over the somatotopically appropriate scalp regions – i.e. contra-lateral following hand stimulation (e.g. compare power change at high beta-gamma between hemisphere contra and ipsi-lateral to stimulus) and midline central channels following feet stimulation (Fig. 1c and d and Fig. 2c and d).

**Figure 2.**
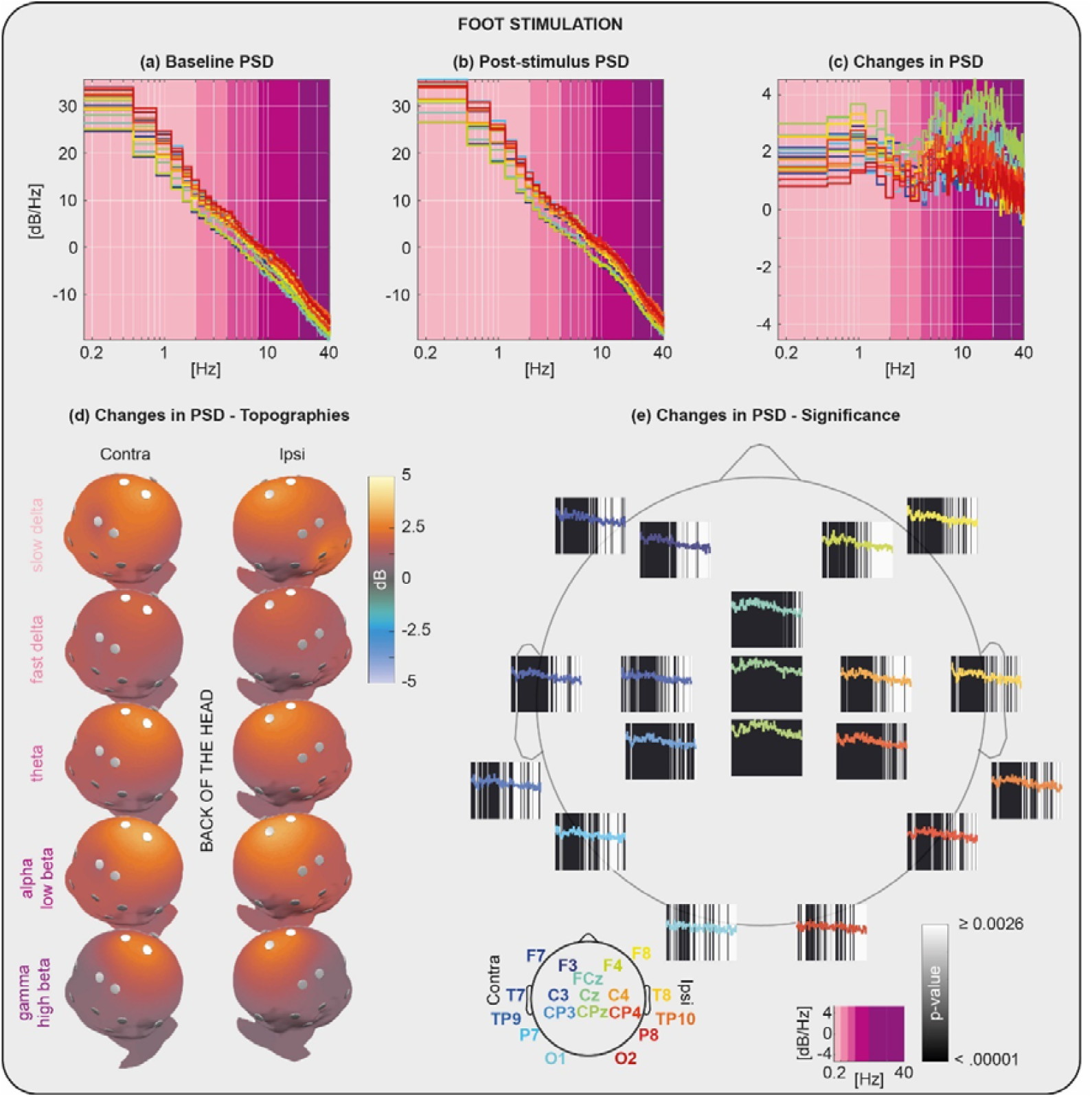
Spectro-spatial EEG power changes associated with tactile stimulation of the feet. Power spectral density (PSD) at baseline **(a)** and following stimulation **(b)** at each channel. PSD of changes elicited by the stimulation **(c)** and 3D maps of their scalp distribution in the slow-delta (0.2-2 Hz), fast-delta (2-4 Hz), theta (4-8 Hz), alpha-low beta (8-20 Hz) and high beta-gamma (20-40 Hz) band **(d)**. Statistical significance of the power changes at each channel **(e)**, note that the changes are the same as those in panel (c) but on a linear frequency axis to allow the display of faster frequencies. P-value limits relate to adjustment for multiple comparisons.

### Spontaneous activity of somatosensory cortical networks dampens EEG power changes evoked by tactile stimulation

We then sought to determine whether the spontaneous activity of somatosensory cortical networks influenced their recruitment by external stimulation. Since we had no control over the ongoing activity at the time of stimulation, we examined whether these networks happened to be spontaneously active just prior to each stimulus. To assess this, we evaluated how baseline activity with spectro-spatial characteristics similar to those evoked by somatosensory stimulation (i.e., a somatosensory-evoked template) influenced the power changes elicited by the stimulus. Specifically, we quantified the similarity between the baseline EEG power distribution and the somatosensory-evoked template for each trial (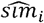), and examined how this similarity related to the EEG power change evoked by the corresponding stimulus (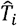) (Fig. 3). Similarity could range between 0-5, where 0 is no similarity between topographies and 5 is an exact match. A close match between the spectro-spatial distribution of baseline power to the somatosensory-evoked template would imply that the somatosensory cortical networks were spontaneously active before the stimulus.

**Figure 3.**
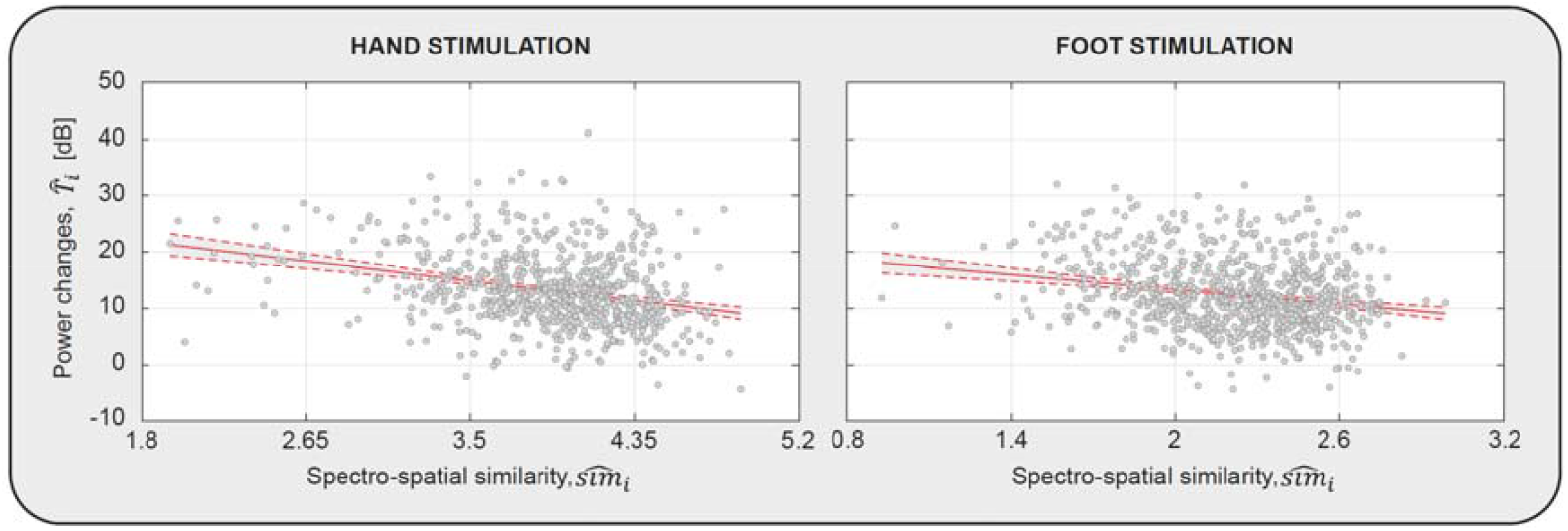
Spontaneous activity influences EEG power changes evoked by tactile stimulation. Scatter plots of evoked power changes (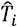) vs spectro-spatial distribution similarity to sensory-evoked activity (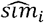) for hands and feet stimulation. The solid red line is the regression fit to data points (dashed red lines are the 95% confidence bounds for the mean). In both panels each data point represents a single trial.

We found that higher spectro-spatial distribution similarity was associated with smaller stimulus-evoked power changes. Power changes were significantly affected by the spontaneous activity preceding the stimulus as demonstrated by the linear model 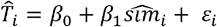, performing better in explaining the variance in 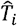 than a constant model (hands: r^2^ = 0.09, f-stats vs constant model = 67, p < 0.0001; feet: r^2^ = 0.05, f-stats vs constant model = 41, p < 0.0001; Fig. 3). Stimulus-evoked power changes at mean similarity were 13.21 dB *(β*_*0*_ hands, t = 56.37, p < 0.0001) and 12.42 dB (*β*_*0*_feet, t = 58.54, p < 0.0001). These stimulus-evoked power changes decreased by 4.1 dB (*β*_*0*_ hands, t = −0.18, p < 0.0001) and 4.3 dB (*β*_*0*_ feet, t = −6.39, p < 0.0001) for every extra 1.0 degree of similarity (range: 1.9-4.7 hands, 1.0-2.9 feet).

## Discussion

We demonstrated that spontaneous cortical activity with spectro-spatial distribution similar to that of somatosensory-evoked activity dampened power changes elicited with stimulation. This suggests that spontaneous activation of focal and extended somatosensory networks depresses their excitability in preterm human neonates. Tactile stimulation elicited an increase in EEG power, but the extent of this increase was dependent on the spectro-spatial characteristics of the activity preceding the stimulus: the closer the spectro-spatial baseline cortical activity was to that evoked by the somatosensory stimulus, the weaker was the change elicited. This result offers an etiological explanation to the discontinuous pattern of cortical activity at this age (burst-quiescence), which may be due to network refractoriness that recovers during inter-burst intervals (Vyazovskiy et al., 2009).

### Tactile stimulation evokes focal and extended cortical activity changes in preterm infants

External tactile (Milh et al., 2007; Vanhatalo et al., 2009; Whitehead et al., 2018b; Leikos et al., 2020), noxious (Fabrizi et al., 2011; Green et al., 2018), visual (Colonnese et al., 2010) and auditory (Chipaux et al., 2013; Kaminska et al., 2018) stimuli elicit rudimentary cortical responses from as early as 28 weeks gestation in humans. These responses are generally composed of fast ripples overriding a larger slow wave focussed around the scalp locations overlaying the primary sensory areas relevant to stimulus modality (i.e. central – tactile; occipital – visual). Here we show that these responses (to tactile stimuli) have complex spectro-spatial characteristics, where oscillations at different frequencies have different voltage fields. The tactile response had a wide band power spectrum but included two clear peaks within the alpha-beta-gamma and slow-delta range and a dip in the theta range. Such coupled oscillatory signatures are a hallmark of activity-dependent processes at time scales relevant for synaptic plasticity (Buzsáki et al., 2013). Changes above 20 Hz (in the beta-gamma range) were localised to central contralateral and midline areas following hand and foot stimulation respectively, while those at the slow end of the spectrum were more widespread.

The results described above could imply analogous mechanisms to those observed in rodent pups, in which somatosensory stimulation elicits distinct patterns of oscillations which differentially synchronise neonatal cortical networks (Yang et al., 2009). Early Gamma Oscillations (EGO) involve focal networks limited to a single cortical column or barrel (Minlebaev et al., 2011; McVea et al., 2012; Yang et al., 2013), while long oscillations extend beyond those boundaries (Yang et al., 2009). EGO are driven by thalamic oscillators and are thought to support the development of thalamocortical projections and drive the organization of vertical topographic functional units (Minlebaev et al., 2011), while long oscillations may serve as a tangential signal binding these units together within the somatomotor area (Yang et al., 2009; Quairiaux et al., 2011; An et al., 2014). As thalamo-cortical connections’ formation precedes that of cortico-cortical connections in the human brain (Volpe, 2009), a role for slow oscillations in synchronising cortico-cortical ensembles may underlie why sensory-evoked delta activity declines later in preterm infants than fast ripples (Chipaux et al., 2013; Whitehead et al., 2018b). Our results suggest that sensory evoked gamma activity may represent the focal activation of columnar functional sensory units, while slow activity may represent horizontal coordination between them within the contralateral somatomotor area (Allievi et al., 2016).

### Spontaneous engagement of somatosensory network suppresses their externally-driven excitability

Cortical bursts in preterm infants can be evoked by sensory stimulation, but can also occur spontaneously in the absence of external stimuli (Whitehead et al., 2016) (Supplementary Video 2). These spontaneous activity patterns occur with varying spatial configurations (Arichi et al., 2017) which overlap with those evoked by external stimulation (Whitehead et al., 2016). Here we show that the response to a somatosensory input is suppressed if the background cortical activity preceding stimulus onset resembles that evoked by the stimulus itself.

In neonatal animal models, spontaneous activity starts from central pattern generators in the sensory periphery, spinal cord, thalamus or subplate (Martini et al., 2021). Independently of their source, when these patterns propagate to the cortex, they ultimately involve the same regions which are engaged by external stimuli, and are confined within the domain related to one sensory modality (e.g. somatosensory: (Mizuno et al., 2018; Nakazawa et al., 2020); visual (Ruthazer and Stryker, 1996; Kenet et al., 2003; O’Hashi et al., 2018; Smith et al., 2018)). Concordant with this, in human preterm infants, resting-state fMRI activity involves the same areas engaged by sensory stimulation, suggesting that these same networks can also be coherently active in the absence of any external stimulus (Doria et al., 2010; Allievi et al., 2016). As specific patterns of cortical activation correspond to specific patterns of voltage field distribution over the scalp (Arichi et al., 2017), background EEG with spectro-spatial characteristics similar to that evoked by somatosensory stimulation is likely to represent the spontaneous unsolicited activation of developing somatosensory networks.

Whether spontaneous or evoked, early activity patterns are supported by a mutually excitatory loop which comprises subplate neurons: a transient neuronal population in the immature brain, projecting to and receiving glutamatergic input from layers 4-6 of the cortical plate (Tolner et al., 2012; Viswanathan et al., 2012). Such a recurrent excitatory circuit will encounter activity-dependent synaptic depression, possibly because of transmitter depletion and ligand-induced changes in postsynaptic currents (Fedirchuk et al., 1999; Tabak et al., 2000). This depletion will transiently reduce network excitability, which then slowly recovers allowing for the next neuronal event, whether spontaneous or evoked, to occur. Our results therefore suggest that the spontaneous activation of somatosensory networks in preterm human neonates has a major dampening impact on the processing of later ascending sensory input.

In conclusion, we offer evidence that sensory-evoked activity in preterm human neonates represents the coordinated activation of local (columnar) and extended (tangential) cortical aggregates and that the occurrence of spontaneous cortical events in the same regions depresses their excitability preventing their immediate re-engagement. This “refractoriness” offers an etiological explanation to the cyclical burst-quiescence pattern typical of preterm cortical activity.

## Acknowledgements

This work was supported by the Medical Research Council UK (MR/L019248/1 and MR/S003207/1) primarily (awardee: LF), Brain Research UK (awardee: KW), and the 7th European Community Framework Programme via Grant CoG 2015-682172NETS (awardee: SO). We would like to thank the families who participated in this research.

## Supplementary Material

Supplementary Video 1: Two examples of tactile stimulation of the left hand in a 35 weeks post-menstrual age infant, during natural sleep. To maintain the infant’s comfort, they remain wrapped in their bedding, with only a small amount of skin uncovered to deliver the tap.

Supplementary Video 2: Example of a spontaneous cortical burst in a 31 weeks post-menstrual age infant, during natural sleep.

## Bibliography

Allievi AG, Arichi T, Tusor N, Kimpton J, Arulkumaran S, Counsell SJ, Edwards AD, Burdet E (2016) Maturation of Sensori-Motor Functional Responses in the Preterm Brain. Cerebral Cortex 26:402–413.

An S, Kilb W, Luhmann HJ (2014) Sensory-Evoked and Spontaneous Gamma and Spindle Bursts in Neonatal Rat Motor Cortex. J Neurosci 34:10870–10883.

André M, Lamblin M-D, d’Allest AM, Curzi-Dascalova L, Moussalli-Salefranque F, Nguyen The Tich S, Vecchierini-Blineau M-F, Wallois F, Walls-Esquivel E, Plouin P (2010) Electroencephalography in premature and full-term infants. Developmental features and glossary. Neurophysiologie Clinique/Clinical Neurophysiology 40:59–124.

Arichi T, Whitehead K, Barone G, Pressler R, Padormo F, Edwards AD, Fabrizi L (2017) Localization of spontaneous bursting neuronal activity in the preterm human brain with simultaneous EEG-fMRI Kastner S, ed. eLife 6:e27814.

Bourel-Ponchel E, Gueden S, Hasaerts D, Héberlé C, Malfilâtre G, Mony L, Vignolo-Diard P, Lamblin M-D (2021) Normal EEG during the neonatal period: maturational aspects from premature to full-term newborns. Neurophysiologie Clinique 51:61–88.

Buzsáki G, Logothetis N, Singer W (2013) Scaling Brain Size, Keeping Timing: Evolutionary Preservation of Brain Rhythms. Neuron 80:751–764.

Chipaux M, Colonnese MT, Mauguen A, Fellous L, Mokhtari M, Lezcano O, Milh M, Dulac O, Chiron C, Khazipov R, Kaminska A (2013) Auditory Stimuli Mimicking Ambient Sounds Drive Temporal “Delta-Brushes” in Premature Infants. PLOS ONE 8:e79028.

Colonnese MT, Kaminska A, Minlebaev M, Milh M, Bloem B, Lescure S, Moriette G, Chiron C, Ben-Ari Y, Khazipov R (2010) A Conserved Switch in Sensory Processing Prepares Developing Neocortex for Vision. Neuron 67:480–498.

Doria V, Beckmann CF, Arichi T, Merchant N, Groppo M, Turkheimer FE, Counsell SJ, Murgasova M, Aljabar P, Nunes RG, Larkman DJ, Rees G, Edwards AD (2010) Emergence of resting state networks in the preterm human brain. PNAS 107:20015–20020.

Dutta S, Iyer KK, Vanhatalo S, Breakspear M, Roberts JA (2023) Mechanisms underlying pathological cortical bursts during metabolic depletion. Nat Commun 14:4792.

Fabrizi L, Slater R, Worley A, Meek J, Boyd S, Olhede S, Fitzgerald M (2011) A Shift in Sensory Processing that Enables the Developing Human Brain to Discriminate Touch from Pain. Current Biology 21:1552–1558.

Fabrizi L, Verriotis M, Williams G, Lee A, Meek J, Olhede S, Fitzgerald M (2016) Encoding of mechanical nociception differs in the adult and infant brain. Sci Rep 6 Available at: http://www.ncbi.nlm.nih.gov/pmc/articles/PMC4921818/ [Accessed July 4, 2016].

Fedirchuk B, Wenner P, Whelan PJ, Ho S, Tabak J, O’Donovan MJ (1999) Spontaneous Network Activity Transiently Depresses Synaptic Transmission in the Embryonic Chick Spinal Cord. J Neurosci 19:2102–2112.

Feller MB, Wellis DP, Stellwagen D, Werblin FS, Shatz CJ (1996) Requirement for Cholinergic Synaptic Transmission in the Propagation of Spontaneous Retinal Waves. Science 272:1182–1187.

Godfrey KB, Swindale NV (2007) Retinal Wave Behavior through Activity-Dependent Refractory Periods. PLOS Computational Biology 3:e245.

Green G, Hartley C, Hoskin A, Duff E, Shriver A, Wilkinson D, Adams E, Rogers R, Moultrie F, Slater R (2018) Behavioural discrimination of noxious stimuli in infants is dependent on brain maturation. PAIN Articles in Press Available at: https://journals.lww.com/pain/Abstract/publishahead/Behavioural_discrimination_of_noxious_stimuli_in.98819.aspx [Accessed November 15, 2018].

Guérit J-M, Amantini A, Amodio P, Andersen KV, Butler S, de Weerd A, Facco E, Fischer C, Hantson P, Jäntti V, Lamblin M-D, Litscher G, Péréon Y (2009) Consensus on the use of neurophysiological tests in the intensive care unit (ICU): Electroencephalogram (EEG), evoked potentials (EP), and electroneuromyography (ENMG). Neurophysiologie Clinique/Clinical Neurophysiology 39:71–83.

Kaminska A, Delattre V, Laschet J, Dubois J, Labidurie M, Duval A, Manresa A, Magny J-F, Hovhannisyan S, Mokhtari M, Ouss L, Boissel A, Hertz-Pannier L, Sintsov M, Minlebaev M, Khazipov R, Chiron C (2018) Cortical Auditory-Evoked Responses in Preterm Neonates: Revisited by Spectral and Temporal Analyses. Cereb Cortex 28:3429–3444.

Kenet T, Bibitchkov D, Tsodyks M, Grinvald A, Arieli A (2003) Spontaneously emerging cortical representations of visual attributes. Nature 425:954–956.

Khazipov R, Luhmann HJ (2006) Early patterns of electrical activity in the developing cerebral cortex of humans and rodents. Trends in Neurosciences 29:414–418.

Kirmse K, Zhang C (2022) Principles of GABAergic signaling in developing cortical network dynamics. Cell Reports 38:110568.

Leikos S, Tokariev A, Koolen N, Nevalainen P, Vanhatalo S (2020) Cortical responses to tactile stimuli in preterm infants. European Journal of Neuroscience 51:1059–1073.

Martini FJ, Guillamón-Vivancos T, Moreno-Juan V, Valdeolmillos M, López-Bendito G (2021) Spontaneous Activity in Developing Thalamic and Cortical Sensory Networks. Neuron 109:2519–2534.

McVea DA, Mohajerani MH, Murphy TH (2012) Voltage-Sensitive Dye Imaging Reveals Dynamic Spatiotemporal Properties of Cortical Activity after Spontaneous Muscle Twitches in the Newborn Rat. J Neurosci 32:10982–10994.

Milh M, Kaminska A, Huon C, Lapillonne A, Ben-Ari Y, Khazipov R (2007) Rapid Cortical Oscillations and Early Motor Activity in Premature Human Neonate. Cerebral Cortex 17:1582–1594.

Minlebaev M, Colonnese M, Tsintsadze T, Sirota A, Khazipov R (2011) Early Gamma Oscillations Synchronize Developing Thalamus and Cortex. Science Available at: <https://www.science.org/doi/abs/10.1126/science.1210574> [Accessed January 26, 2022].

Mizuno H, Ikezoe K, Nakazawa S, Sato T, Kitamura K, Iwasato T (2018) Patchwork-Type Spontaneous Activity in Neonatal Barrel Cortex Layer 4 Transmitted via Thalamocortical Projections. Cell Reports 22:123–135.

Molnár Z, Luhmann HJ, Kanold PO (2020) Transient cortical circuits match spontaneous and sensory-driven activity during development. Science 370 Available at: https://science.sciencemag.org/content/370/6514/eabb2153 [Accessed October 20, 2020].

Nakazawa S, Yoshimura Y, Takagi M, Mizuno H, Iwasato T (2020) Developmental Phase Transitions in Spatial Organization of Spontaneous Activity in Postnatal Barrel Cortex Layer 4. J Neurosci 40:7637–7650.

O’Donovan MJ (1999) The origin of spontaneous activity in developing networks of the vertebrate nervous system. Current Opinion in Neurobiology 9:94–104.

O’Hashi K, Fekete T, Deneux T, Hildesheim R, van Leeuwen C, Grinvald A (2018) Interhemispheric Synchrony of Spontaneous Cortical States at the Cortical Column Level. Cerebral Cortex 28:1794–1807.

Quairiaux C, Megevand P, Kiss JZ, Michel CM (2011) Functional Development of Large-Scale Sensorimotor Cortical Networks in the Brain. Journal of Neuroscience 31:9574–9584.

Ruthazer ES, Stryker MP (1996) The Role of Activity in the Development of Long-Range Horizontal Connections in Area 17 of the Ferret. J Neurosci 16:7253–7269.

Saby JN, Meltzoff AN, Marshall PJ (2016) Beyond the N1: A review of late somatosensory evoked responses in human infants. International Journal of Psychophysiology 110:146–152.

Smith GB, Hein B, Whitney DE, Fitzpatrick D, Kaschube M (2018) Distributed network interactions and their emergence in developing neocortex. Nat Neurosci 21:1600–1608.

Tabak J, Senn W, O’Donovan MJ, Rinzel J (2000) Modeling of Spontaneous Activity in Developing Spinal Cord Using Activity-Dependent Depression in an Excitatory Network. J Neurosci 20:3041–3056.

Tiriac A, Del Rio-Bermudez C, Blumberg MS (2014) Self-Generated Movements with “Unexpected” Sensory Consequences. Current Biology 24:2136–2141.

Tolner EA, Sheikh A, Yukin AY, Kaila K, Kanold PO (2012) Subplate Neurons Promote Spindle Bursts and Thalamocortical Patterning in the Neonatal Rat Somatosensory Cortex. J Neurosci 32:692–702.

Tsuchida TN, Wusthoff CJ, Shellhaas RA, Abend NS, Hahn CD, Sullivan JE, Nguyen S, Weinstein S, Scher MS, Riviello JJ, Clancy RR, American Clinical Neurophysiology Society Critical Care Monitoring Committee (2013) American clinical neurophysiology society standardized EEG terminology and categorization for the description of continuous EEG monitoring in neonates: report of the American Clinical Neurophysiology Society critical care monitoring committee. J Clin Neurophysiol 30:161–173.

Vanhatalo S, Jousmäki V, Andersson S, Metsäranta M (2009) An Easy and Practical Method for Routine, Bedside Testing of Somatosensory Systems in Extremely Low Birth Weight Infants. Pediatr Res 66:710–713.

Viswanathan S, Bandyopadhyay S, Kao JPY, Kanold PO (2012) Changing Microcircuits in the Subplate of the Developing Cortex. J Neurosci 32:1589–1601.

Volpe JJ (2009) The Encephalopathy of Prematurity—Brain Injury and Impaired Brain Development Inextricably Intertwined. Seminars in Pediatric Neurology 16:167–178.

Vyazovskiy VV, Faraguna U, Cirelli C, Tononi G (2009) Triggering Slow Waves During NREM Sleep in the Rat by Intracortical Electrical Stimulation: Effects of Sleep/Wake History and Background Activity. Journal of Neurophysiology 101:1921–1931.

Wallois F, Routier L, Heberlé C, Mahmoudzadeh M, Bourel-Ponchel E, Moghimi S (2021) Back to basics: the neuronal substrates and mechanisms that underlie the electroencephalogram in premature neonates. Neurophysiologie Clinique 51:5–33.

Watanabe K, Hayakawa F, Okumura A (1999) Neonatal EEG: a powerful tool in the assessment of brain damage in preterm infants. Brain and Development 21:361–372.

Whitehead K, Fabrizi L (2024) Sleep-wake resting EEG epochs in human neonates. Available at: https://osf.io/.

Whitehead K, Laudiano-Dray MP, Meek J, Fabrizi L (2018a) Emergence of mature cortical activity in wakefulness and sleep in healthy preterm and full-term infants. Sleep 41 Available at: https://academic.oup.com/sleep/article/41/8/zsy096/4995737 [Accessed August 14, 2018].

Whitehead K, Meek J, Fabrizi L (2018b) Developmental trajectory of movement-related cortical oscillations during active sleep in a cross-sectional cohort of pre-term and full-term human infants. Scientific Reports 8:17516.

Whitehead K, Pressler R, Fabrizi L (2016) Characteristics and clinical significance of delta brushes in the EEG of premature infants. Clinical Neurophysiology Practice 2:12–18.

Worley A, Fabrizi L, Boyd S, Slater R (2012) Multi-modal pain measurements in infants. Journal of Neuroscience Methods 205:252–257.

Yang J-W, An S, Sun J-J, Reyes-Puerta V, Kindler J, Berger T, Kilb W, Luhmann HJ (2013) Thalamic Network Oscillations Synchronize Ontogenetic Columns in the Newborn Rat Barrel Cortex. Cerebral Cortex 23:1299–1316.

Yang J-W, Hanganu-Opatz IL, Sun J-J, Luhmann HJ (2009) Three Patterns of Oscillatory Activity Differentially Synchronize Developing Neocortical Networks In Vivo. J Neurosci 29:9011– 9025.

Zheng J, Lee S, Zhou ZJ (2006) A transient network of intrinsically bursting starburst cells underlies the generation of retinal waves. Nat Neurosci 9:363–371.

